# Development of GPC2-directed chimeric antigen receptors using mRNA for pediatric brain tumors

**DOI:** 10.1101/2021.07.06.451385

**Authors:** Jessica B. Foster, Crystal Griffin, Jo Lynne Rokita, Allison Stern, Cameron Brimley, Komal Rathi, Maria V. Lane, Samantha N. Buongervino, Tiffany Smith, Peter J. Madsen, Daniel Martinez, Robert J. Wechsler-Reya, Katalin Karikó, Phillip B. Storm, David M. Barrett, Adam C. Resnick, John M. Maris, Kristopher R. Bosse

## Abstract

Pediatric brain tumors are the leading cause of cancer death in children with an urgent need for innovative therapies. Here we show that the cell surface oncoprotein glypican 2 (GPC2) is highly expressed on multiple lethal pediatric brain tumors, including medulloblastomas, embryonal tumors with multi-layered rosettes, other CNS embryonal tumors, as well as definable subsets of highly malignant gliomas. To target GPC2 on these pediatric brain tumors with adoptive cellular therapies and mitigate potential inflammatory neurotoxicity, we developed four mRNA chimeric antigen receptor (CAR) T cell constructs using the highly GPC2-specific fully human D3 single chain variable fragment. First, we validated and prioritized these CARs using *in vitro* cytotoxicity and T cell degranulation assays with GPC2-expressing neuroblastoma cells. Next, we expanded the testing of the two most potent GPC2-directed CAR constructs prioritized from these studies to GPC2-expressing medulloblastoma and high-grade glioma cell lines, showing significant GPC2-specific cell death in multiple models. Finally, locoregional delivery bi-weekly of two to four million GPC2-directed mRNA CAR T cells induced significant and sustained tumor regression in two orthotopic medulloblastoma models, and significantly prolonged survival in an aggressive orthotopic thalamic diffuse midline glioma model. No GPC2-directed CAR T cell related neurologic or systemic toxicity was observed. Taken together, these data show that GPC2 is a highly differentially expressed cell surface protein on multiple malignant pediatric brain tumors that can be targeted safely with local delivery of mRNA CAR T cells.

**One Sentence Summary:** Glypican 2 is expressed on the surface of multiple pediatric brain tumors and can be successfully targeted with mRNA chimeric antigen receptor T cells.

## INTRODUCTION

Immunotherapy has developed into a major pillar of cancer therapy over the past two decades(*1*). Chimeric antigen receptor (CAR) T cells are one form of immunotherapy, where targeted T cell killing is achieved through combining the specificity of an antibody with the stimulatory signals of cytotoxic T cells(*2*). CAR T cell therapy has provided dramatic success in leukemias and lymphomas(*3–5*), and tisagenlecleucel recently became the first FDA-approved cellular therapy product(*6*). This achievement has prompted similar investigations for solid tumors but without significant success to date(*7*). Developing effective CAR T cell therapy for solid tumors has many challenges related to difficulty finding appropriate target antigens, the suppressive tumor microenvironment, T cell dysfunction, and poor CAR T cell trafficking(*7*). Among brain tumors, CAR T cell therapy remains in its infancy, with a handful of adult clinical trials showing minimal to transient success(*8–10*), and trials in pediatric brain tumors only just beginning(*11*).

The majority of CAR T cell work to date has used viral vectors to produce constitutively expressed CAR molecules(*12*). However, CAR molecules may also be introduced into T cells using *in vitro* transcribed mRNA(*13*), and we have shown effective CAR T cells can be made utilizing the same innovative technology that recently allowed mRNA to be successfully employed for COVID-19 vaccination development(*14, 15*). Using mRNA to generate CAR T cells for adoptive transfer may offer several advantages in the clinic or in preclinical validation of CARs. First, mRNA transfection is a more efficient process than viral transduction, with uniformly high levels of CAR on the surface(*16*). Second, due to the quicker production time, iterative changes in the CAR binding site or structure are more easily tested(*17*). Third, mRNA is less expensive than viral vectors when scaling up to clinical grade application. Finally, when exploring novel immunotherapeutic targets, transient expression may be desirable to ensure safety, particularly in the brain where off-target toxicity may cause significant morbidity and mortality(*18, 19*). The major disadvantage of using mRNA CAR T cells is the transient nature, which may not offer the persistence needed for full eradication of disease(*17*), but this could be addressed by repetitive dosing of a fully human CAR construct. Prior work has shown that while systemic delivery of mRNA CAR T cells does not result in long term disease control(*20–22*), local delivery into solid tumors can be remarkably effective(*23, 24*).

We have recently shown that Glypican 2 (GPC2) is a heparan-sulfate proteoglycan cell-surface oncoprotein that is overexpressed on the surface of neuroblastomas as well as several other pediatric and adult cancers(*25, 26*). GPC2 appears to harbor all the qualities of a *bona fide* immunotherapeutic target, including cell-surface location, tumor-specific expression, tumor dependence and enrichment in the tumor stem cell compartment(*25, 26*). To therapeutically capitalize on this differential expression, we and others have identified several GPC2-specific binders(*25, 27, 28*) and shown robust efficacy of GPC2-targeting antibody-drug conjugates (ADCs)(*25, 26*), immunotoxins(*27*), and DNA vector-derived CAR T cells(*27, 28*). Here we sought to build upon these data with a specific focus on GPC2 in pediatric brain cancers, including better defining the brain tumor histotypes that harbor high-levels of GPC2 expression, the genomic biomarkers correlated with GPC2 expression, and in creating GPC2-redirected mRNA CAR T cells to enable a safe and efficacious therapeutic approach for local intracranial delivery.

## RESULTS

### GPC2 is expressed in multiple pediatric brain tumors and correlated with MYCN expression and chromosome 7q gain

GPC2 is a differentially expressed MYCN-regulated oncoprotein found on the surface of most neuroblastomas and several other embryonal cancers, including medulloblastomas and retinoblastomas(*25, 26*). To better define *GPC2* expression specifically in pediatric brain tumors, we analyzed RNA sequencing data from 20 unique pediatric CNS tumor types (total n=833) in the Pediatric Brain Tumor Atlas (OpenPBTA), which revealed that several additional pediatric CNS tumors harbor high levels of *GPC2* RNA (**Fig 1A**). Based on normal expression data from the Genotype-Tissue expression (GTex) portal, we chose a cutoff value of a TPM of 10 to delineate high *GPC2* expression. Evaluation of tumors throughout the OpenPBTA biorepository showed high *GPC2* expression in 100% of embryonal tumors with multilayered rosettes (ETMR), 95% of medulloblastomas, 93% other CNS embryonal tumors (CNS embryonal tumor, NOS), 50% of choroid plexus carcinomas (CPC), 41% of high-grade gliomas (HGG) without H3 K28M mutation, and 40% of diffuse midline gliomas (DMG) (**Fig 1B**). While several tumor histotypes were uniform in their *GPC2* expression levels, multiple subtypes had significant heterogeneity in *GPC2* expression motivating us to look further at *GPC2* expression across tumor molecular subgroups where possible. First, for medulloblastomas, we observed the highest *GPC2* expression in group 4 compared to the other subtypes (Wnt, SHH, and group 3; p<0.01) (**Fig 1C, left**), confirming our prior results(*25*). Across high grade glioma subgroups, H3 G35 mutant gliomas had the highest *GPC2* expression (p<0.05; **Fig 1C, middle**). Embryonal tumors showed high expression across all subgroups (**Fig 1C, right**).

**Fig. 1.**
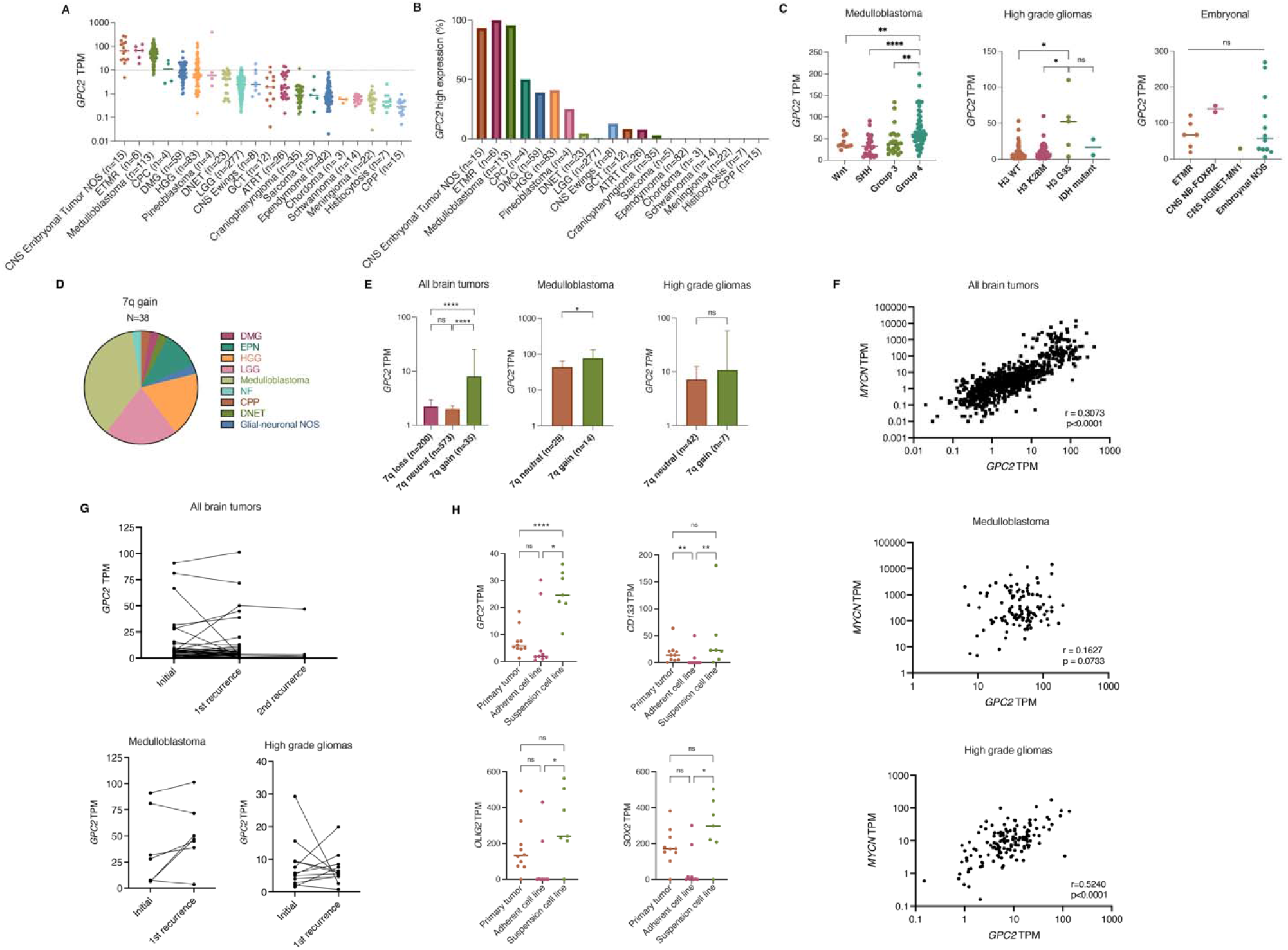
*GPC2* is expressed in pediatric brain tumors. (**A**) *GPC2* RNA sequencing data across OpenPBTA pediatric brain tumors cohorts. (**B**) Using a cut-off value of 10 TPM, percentage of tumors in OpenPBTA dataset with high expression of *GPC2*. (**C**) RNA sequencing of *GPC2* expression in medulloblastoma, high grade glioma, and embryonal tumor subtypes. (**D**) Summary of tumor histotypes with chromosome 7q gain. (**E**) *GPC2* expression stratified by chromosome 7q status. (**F**) Correlation of *MYCN* and *GPC2* expression across all pediatric brain tumor samples (top), medulloblastomas (middle), and high-grade gliomas (bottom). (**G**) *GPC2* expression at primary diagnosis and recurrence across all pediatric brain tumors (top), medulloblastomas (bottom left), and high-grade gliomas (bottom right). (**H**) *GPC2*, *CD133*, *SOX2,* and *OLIG2* expression from primary tumor-derived adherent (FBS) or suspension cell lines (serum free media). TPM = transcripts per million, NOS = not otherwise specified, ETMR = embryonal tumor with multilayered rosettes, CPC = choroid plexus carcinoma, DMG = diffuse midline glioma, HGG = high grade glioma, LGG = low grade glioma, DNET = dysembryoplastic neuroepithelial tumor, ATRT = atypical teratoid rhabdoid tumor, GCT = germ cell tumor, and CPP = choroid plexus papilloma. Individual cases indicated by dots, median indicated by line in **A, C** and **H**. Data in **E** displayed as median with 95% confidence interval. ****, p<0.0001; **, p<0.01; *, p<0.05; ns, not significant.

We have also previously found that significantly higher levels of *GPC2* expression are found in neuroblastoma or medulloblastoma tumors that harbor either high level expression of a MYC gene family member and/or contain somatic gain of the *GPC2* locus at chromosome 7q22(*25*). To determine if similar drivers of *GPC2* expression are present across this diverse set of pediatric brain tumors we stratified tumors by chromosome 7q copy number and found that 6.7% of tumors had 7q gain (38 of 565 tumors with copy number data available). The majority of those were medulloblastoma (n=14), LGG (n=8) and HGG (n=7) (**Fig 1D**). Those tumors with 7q gain showed significantly higher *GPC2* expression (p<0.0001; **Fig 1E, left**). This chromosome 7q gain and increased *GPC2* expression correlation was also found in medulloblastomas (p<0.05) **(Fig 1E, center)**, and the same trend was found in high grade gliomas (**Fig 1E, right**). Furthermore, we also observed a positive correlation between *GPC2* and *MYCN* expression within this pediatric brain tumor dataset (r=0.3073, p<0.0001; **Fig 1F, top**), suggesting that MYCN may also regulate *GPC2* expression in some pediatric brain tumors as well. This *MYCN/GPC2* expression correlation was most robust in high grade gliomas (r=0.524, p<0.0001; **Fig 1F, bottom**), but this correlation was not found in medulloblastoma (r=0.16, p=ns; **Fig 1F, middle**). *MYC* expression also showed a weak positive correlation with *GPC2* expression across all brain tumors (r=0.21779, p<0.0001), and combined *MYCN* and *MYC* expression for all tumors was also correlated with higher *GPC2* expression (r=0.3351, p<0.0001).

Next, to evaluate if *GPC2* expression is maintained at the time of brain tumor relapse, we evaluated 47 unique patients with paired diagnosis-relapse tumor RNA sequencing data, including 7 patients with multiple relapsed tumor specimens (**Fig 1G**). *GPC2* expression was generally similar at the time of diagnosis and relapses across all brain tumor histotypes. Looking specifically at medulloblastomas and HGGs, again *GPC2* expression was generally stable, however with individual cases showing both increases and decreases in *GPC2* expression at the time of relapse (**Fig 1G, bottom**).

OpenPBTA data generation is completed on patient samples provided through the Children’s Brain Tumor Network (CBTN). The CBTN also performs cell line generation both in serum-free media and media containing fetal bovine serum (FBS). Cell lines have been generated for ten HGG primary tumors in both media types, with subsequent transcriptome profiling with RNA sequencing, providing the opportunity to compare *GPC2* expression in these paired *in vitro* models(*29*). *GPC2* expression was increased in HGG cell lines generated in serum-free conditions (p<0.01; **Fig 1H**). This increase in *GPC2* expression mirrored the expression trends of other neuronal stem cell markers Olig2, SOX2, and CD133 (**Fig 1H**), suggesting that serum-free conditions better propagate stem-like cells from tumors(*30, 31*) and that *GPC2* expression is increased in the HGG stem cell compartment, similar to what we have previously described in neuroblastomas and small cell lung cancers(*26*).

Finally, we next looked to validate these RNA data by quantifying GPC2 expression in a subset of pediatric brain tumors using immunohistochemistry (IHC), flow cytometry, and western blot. First, we evaluated GPC2 expression by IHC, utilizing a HGG-focused tumor microarray which includes 73 HGGs (glioblastomas, anaplastic astrocytomas, and DMGs), as well as 6 CNS embryonal tumors NOS, 4 medulloblastomas, 3 pineoblastomas, 2 ETMRs, 2 ependymomas, 2 ATRTs, 1 high-grade neuroepithelial tumor with BCOR alteration (HGNET-BCOR), 1 oligodendroglioma, and 1 LGG. Most of the HGG tumors harbored positive GPC2 staining (**Fig 2A**), with very strong staining in the single H3 G35 mutant tumor included in the array (**Fig 2B**). In addition, strong GPC2 staining was also observed in 2 out of 4 medulloblastomas, 1 of 2 ETMRs, 2 of 3 pineoblastomas, 1 of 1 HGNET-BCOR, and 2 of 6 embryonal tumors NOS (**Fig 2B**). GPC2 flow cytometry of medulloblastoma (n= 4) and HGG (n=6) cell lines confirmed robust GPC2 cell-surface expression comparable to the neuroblastoma cell line SMS-SAN (**Fig 2C, D).** Of note the two cell lines with the highest GPC2 cell surface expression were a group 4 medulloblastoma (7316-4509) and a H3 G35 HGG (7316-158), consistent with transcriptomic profiling and IHC. Finally, total GPC2 protein expression in medulloblastoma and HGG cell lines by Western blotting was also found to be comparable to the SMS-SAN neuroblastoma cell line (**Fig 2E**). Taken together, these data show that GPC2 is highly expressed on a cohort of malignant pediatric brain tumors, supporting the testing of GPC2 immune-based therapies in these lethal childhood CNS tumors.

**Fig. 2.**
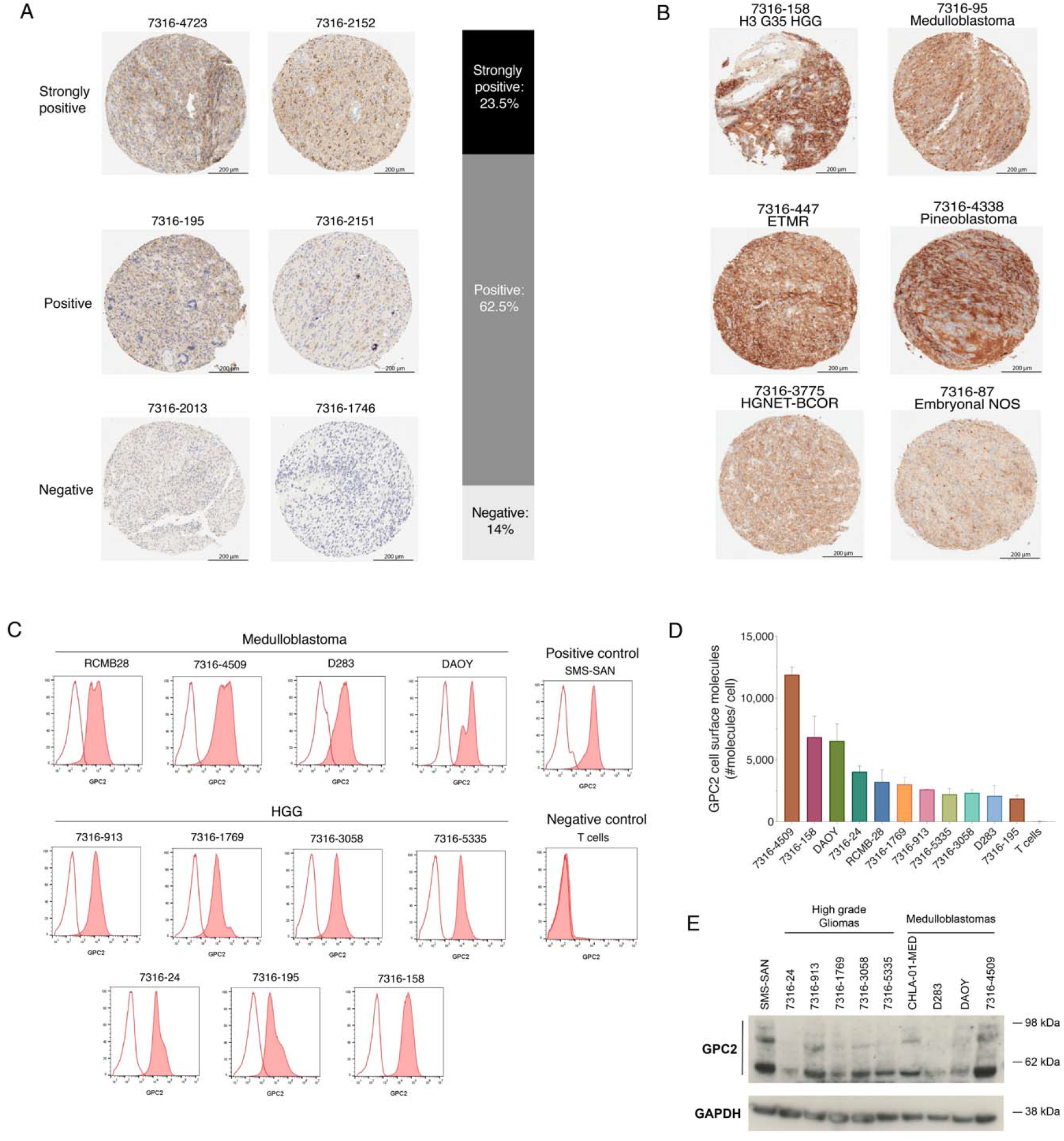
GPC2 protein is expressed on the surface of pediatric brain tumors. (**A**) GPC2 IHC of primary HGGs (n=67) with representative GPC2 IHC staining examples (left) and GPC2 IHC summary (right). (**B**) GPC2 IHC showing strongly positive staining in additional brain tumor histotypes including H3 G35 mutant HGG, medulloblastoma, ETMR, pineoblastoma, HGNET-BCOR, and CNS embryonal tumor NOS. (**C**) GPC2 flow cytometry representative histograms of HGG and medulloblastoma cell lines. Red shaded curves indicate D3-GPC2-IgG1-PE staining, non-shaded curves indicate isotype control. SMS-SAN shown as neuroblastoma positive control and T cells as negative control. (**D**) Summary of the relative quantification of GPC2 molecules per cell from flow cytometry of HGG and medulloblastoma cell lines (n=11). Data is represented as mean ± SEM of 3 independent experiments. (**E**) GPC2 Western blot of a panel of medulloblastoma and HGG cell lines (n=9). SMS-SAN also presented as neuroblastoma positive control. HGG, high grade glioma; ETMR, embryonal tumor with multilayered rosettes; HGNET-BCOR, high grade neuroepithelial tumor with BCOR alteration.

### mRNA GPC2 CAR T cells are cytotoxic to GPC2-expressing neuroblastoma and brain tumor cell lines

For GPC2 CAR generation, we used the single chain variable fragment (scFv) portion of the D3 GPC2 antibody, which binds a conformational epitope of the GPC2 extracellular domain that is identical between human and mouse(*25, 26*). We explored four different configurations of the D3 scFv by manipulating the orientation of the heavy and light chains, as well as the glycine-serine linker length between the heavy and light chains (5 versus 15 amino acids), to identify the structure that provided optimal CAR expression, cell killing, and T cell activation (**Fig 3A)**. All four constructs included a CD8 hinge and transmembrane domain with 41BB costimulatory domain and CD3ζ signaling. CAR mRNA was *in vitro* transcribed and transfected into human T cells, and after 24 hours we evaluated CAR expression and binding to GPC2 via incubation with GPC2-PE. All four constructs showed GPC2 specific binding compared to GD2-directed CAR control (mean MFI 18,721 to 54,849 for constructs versus 684 for control, p<0.001) (**Fig 3B)**. Serial GPC2 CAR expression as measured by protein L flow cytometry showed that CAR expression typically persisted for 5-7 days (**Fig 3C**). To investigate T cell exhaustion, we also measured the expression of negative checkpoint regulators on the surface of GPC2 CAR transfected T cells and found no difference in checkpoint molecule expression across the 4 different constructs (**Fig 3D**).

**Fig. 3.**
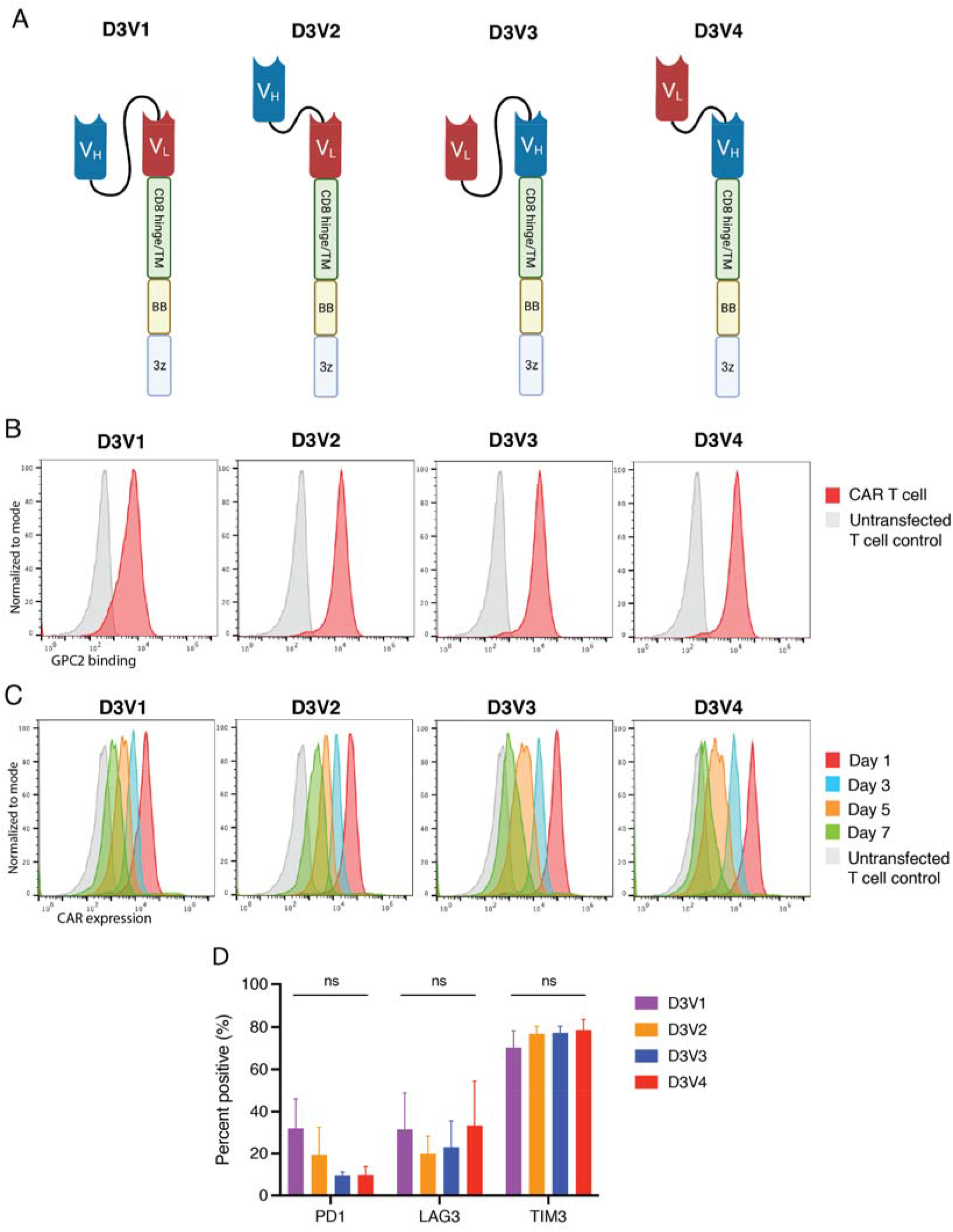
D3-binder based GPC2-directed mRNA CARs specifically bind GPC2 and are transiently expressed on T cells. (**A**) Schematic of GPC2 CAR constructs. (**B**) Flow cytometry representative histograms of GPC2 specific binding by GPC2 CAR T cell constructs. Red line is CAR T cell, grey line is untransfected T cell control. (**C**) Flow cytometry representative histograms of GPC2 CAR persistence on T cells. Red line represents day 1 post-transfection, blue line represents day 3, orange line day 5, and green line day 7. Grey line is untransfected T cell control. (**D**) Cell surface expression of negative checkpoint regulators PD1, Lag3 and Tim3 quantified with flow cytometry at day 4 post-transfection. Data is represented as mean ± SEM of three independent experiments. V_H_, variable heavy chain; V_L_, variable light chain; T_M_, transmembrane domain; BB, 41-BB co-stimulatory domain; 3z, CD3ζ co-stimulatory domain; ns, not significant. Graphics in **A** created with BioRender.com.

We next evaluated the cytotoxicity of CAR T cell constructs *in vitro* using multiple well-characterized neuroblastoma cell lines with variable levels of GPC2 cell surface expression(*25*). CAR T cells transfected with all 4 GPC2 CAR constructs were able to exert robust killing of the endogenously GPC2-high cell lines SMS-SAN and Nb-EbC1 (p<0.0001) (**Fig 4A**). Limited killing was observed of GPC2 low expressing native SKNAS and control 293T cells (10:1 E:T; **Fig 4B**), with robust killing restored with forced overexpression of GPC2 in SKNAS (p<0.001) (**Fig 4C**). At lower E:T ratios (5:1) there was differential cytotoxicity between GPC2 CARs with the D3V3 and D3V4 constructs performing superiorly, most notable on the NB-EbC1 cell line (**Fig 4A**). Furthermore, there was differential T cell activation across the 4 GPC2 CARs, with the D3V3 and D3V4 constructs showing the highest levels of interferon gamma (IFNγ) and interleukin 2 (IL-2) release by ELISA upon co-incubation with GPC2-high neuroblastoma cells (p<0.0001) (**Fig 4D**). Considering these data, the D3V3 and D3V4 GPC2 CAR constructs were prioritized for further testing in brain tumor models based on their superior *in vitro* cytotoxicity and T cell activation.

**Fig. 4.**
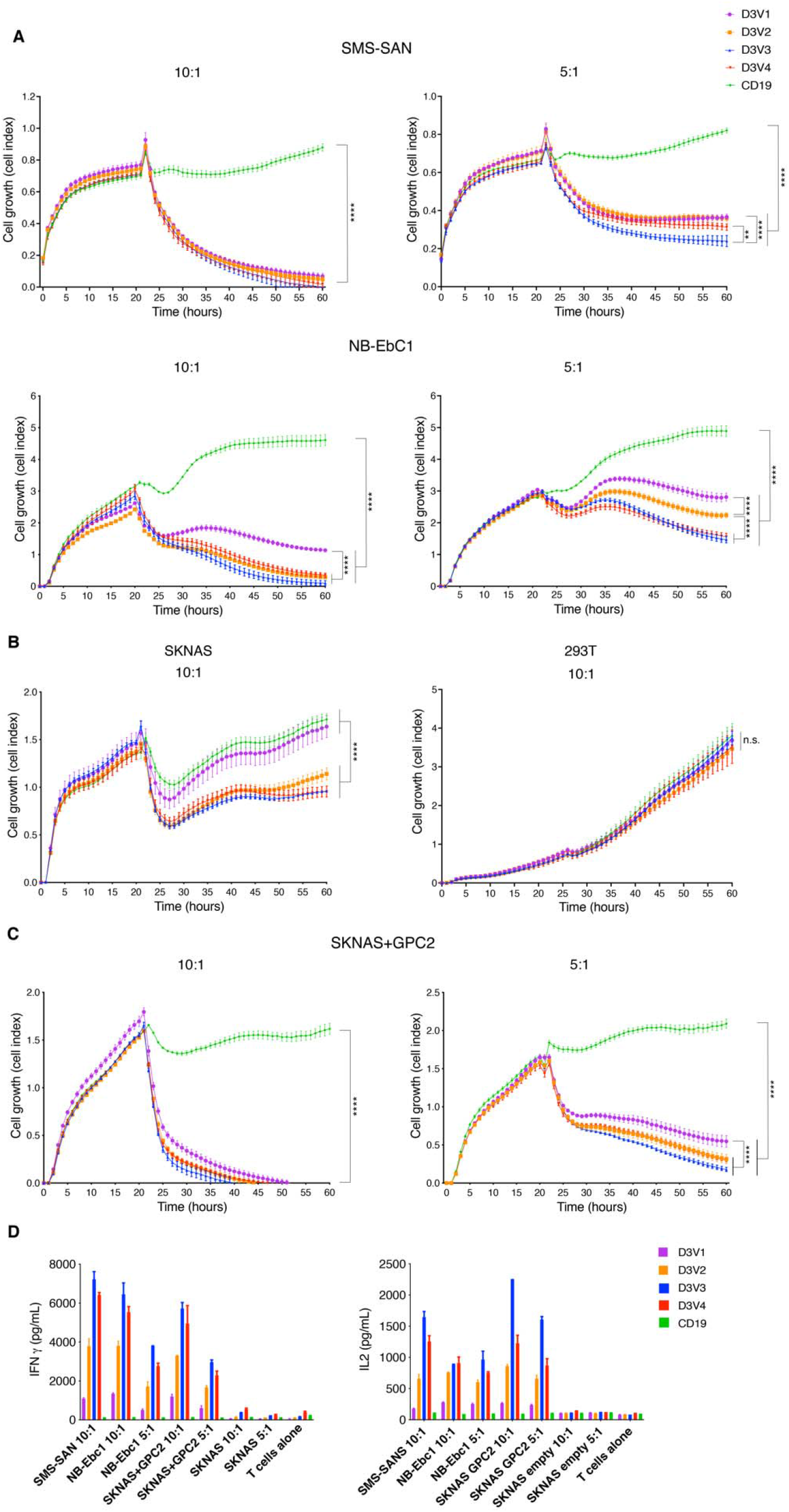
GPC2-directed CAR T cells produce robust cytotoxicity in neuroblastoma cell line models. (**A-C**) GPC2-directed CAR T cells co-incubated on RT-CES with endogenously high (A) and low (B) GPC2 expressing neuroblastoma cell lines, and cell line with forced overexpression of GPC2 (C). (**D**) Cytokine expression measured 24 hours after co-incubation of neuroblastoma and GPC2-directed CAR T cells. E:T, Effector to target ratio. Data is shown as mean ± SEM from a representative experiment with each experiment being done two to four independent times. ****, p<0.0001; **, p<0.01; ns, not significant.

We next tested the cytotoxicity of the prioritized D3V3 and D3V4 GPC2-directed CARs in a panel of medulloblastoma and HGG cell lines with different levels of GPC2 expression (**Fig 2D**), using real-time cell electronic sensing (RT-CES) assay for adherent cell lines (DAOY, 7316-913, and 7316-5335) and luciferase assays for suspension cell lines (7316-4509, 7316-3058, and 7316-24). GPC2-expressing brain tumor cells were selectively killed with GPC2 CAR T cells, but not the CD19 control CAR T cells at E:T ratios of 5:1 or higher (p<0.0001) (**Fig 5A and 5B**). GPC2-directed CAR T cells also showed significant IFNγ release in the presence of GPC2-expressing brain tumor cell lines compared to CD19-directed CAR T cells (p<0.0001) (**Fig 5C**).

**Fig. 5.**
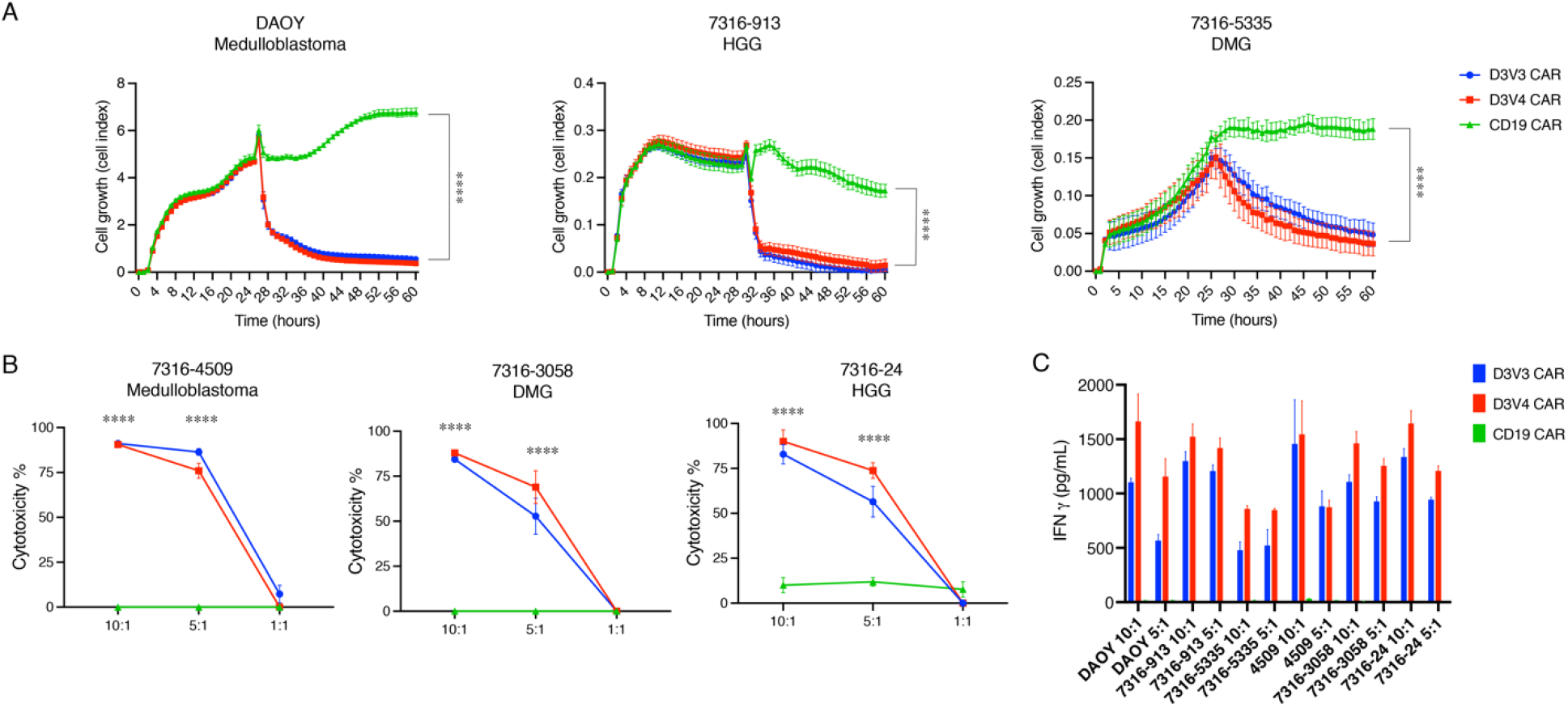
GPC2-directed CAR T cells produce robust cytotoxicity in pediatric brain tumor cell line models. (**A**) GPC2-directed CAR T cells co-incubated with medulloblastoma (DAOY), HGG (7316-913), and DMG (7316-5335) cell lines. E:T ratio is 5:1. (**B**) GPC2 CAR T cells co-inbuated with medulloblastoma (7316-4509), HGG (7316-24), and DMG (7316-3058) cell lines with cytotoxicity measured using a luminescence assay at 48 hours. (**C**) Cytokine levels measured 24 hours after co-incubation of GPC2 CAR T cells with each cell line. Data is shown as mean ± SEM from a representative experiment with each experiment being done two to four independent times. ****, p<0.0001.

### mRNA GPC2-redirected CAR T cells induce sustained tumor regression and increased survival in pediatric brain tumor xenograft models

To evaluate the efficacy of the D3V3 and D3V4 mRNA GPC2-directed CAR T cells *in vivo*, we utilized medulloblastoma and DMG xenograft models. Given the shortened duration of activity with mRNA CAR T cells, and recent work on CAR T cells for CNS tumors reporting increased efficacy with locoregional delivery(*9, 32*), we focused on intratumoral delivery of the prioritized D3V3 and D3V4 GPC2 mRNA CAR T cells. Following orthotopic tumor engraftment of 7316-4509 group 4 medulloblastoma in mice, we placed a fixed guide cannula into the tumor bed that would enable serial CAR T cell dosing, and after 14 days with tumor confirmed on bioluminescent imaging we treated mice with 4×10^6^ of either GPC2- or CD19-directed CAR T cells via catheter into the tumor bed twice a week for 2 weeks for a total of 4 doses. Both D3V3 and D3V4 CAR T cell constructs resulted in sustained 7316-4509 medulloblastoma tumor regression (p<0.0001; **Fig 6A**). We next treated the group 3 medulloblastoma xenograft RCMB28(*33*) in an identical manner with 4×10^6^ of either D3V3 or D3V4 GPC2-directed or CD19-directed CAR T cells via catheter into the tumor bed twice a week for a total of 5 doses, which again showed sustained tumor control via bioluminescent imaging (p<0.0001; **Fig 6B**). For both models, tumor regression was also confirmed by IHC at study endpoint (**Fig 6C**). Notably, no evidence of clinical toxicity was observed in these animals other than expected xenogeneic graft-versus-host disease (GVHD)(*34*) seen in both controls and GPC2-directed treatment groups, including no neurologic toxicity, utilizing the CARs engineered from the D3 GPC2 antibody which binds equally the murine and human GPC2 protein(*26*).

**Fig. 6.**
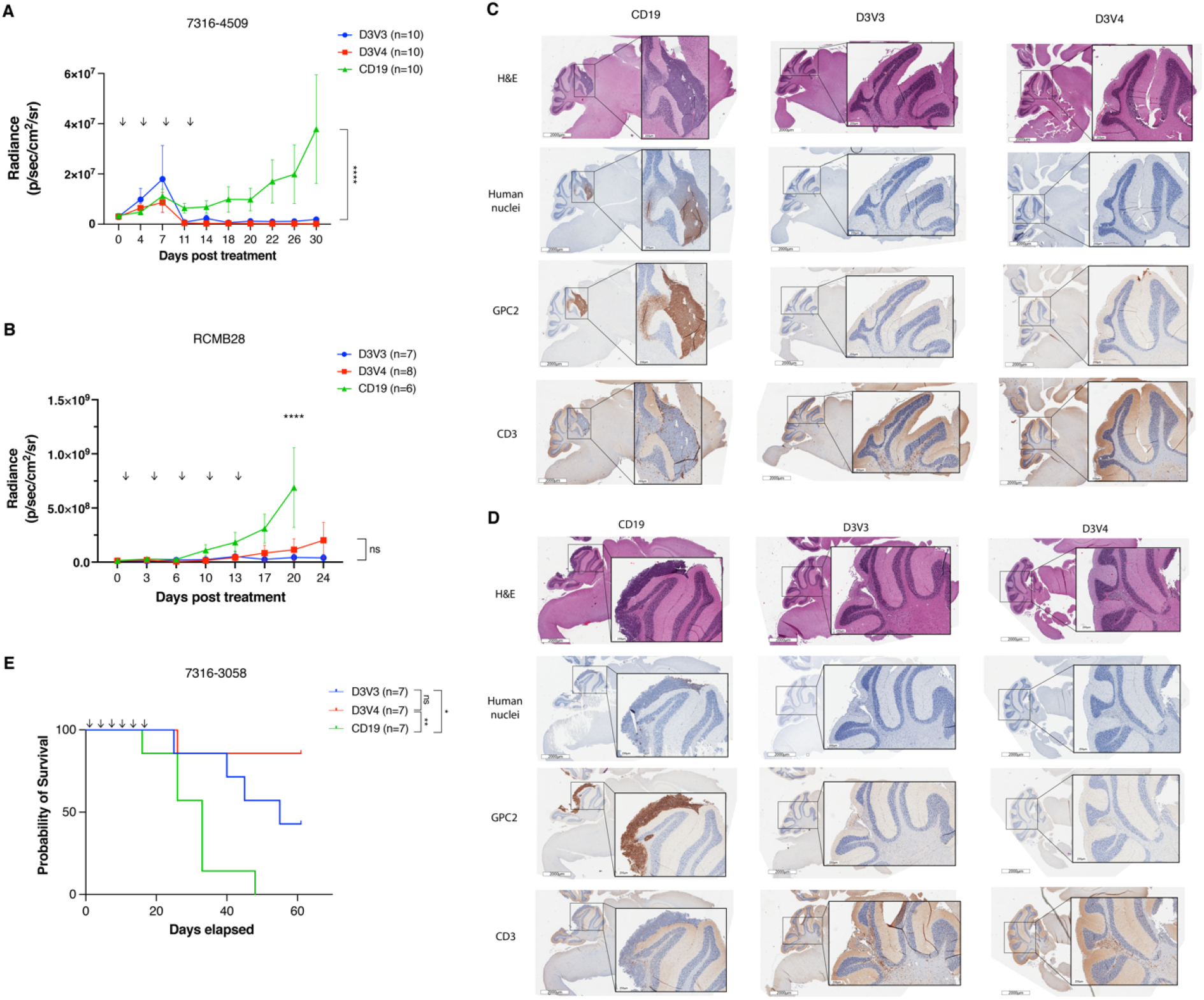
GPC2-directed CAR T cells mediate antitumor responses and prolong survival in pediatric brain tumors *in vivo*. (**A**) Quantification of bioluminescence of the orthotopic group 4 medulloblastoma xenograft 7316-4509 treated with either GPC2 or CD19-directed mRNA CAR T cells. Doses indicated by arrows on graph. Data displayed as mean ± SEM, n= 10 mice per arm. (**B**) Quantification of bioluminescence of the orthotopic group 3 medulloblastoma xenograft RCMB28 treated with either GPC2 or CD19-directed mRNA CAR T cells. Doses indicated by arrows on graph. Data displayed as mean ± SEM, n= 6-8 mice per arm. (**C, D**) IHC of murine brains of a representative mouse per group collected at study endpoint from medulloblastoma xenograft experiments using models 7316-4509 (C) and RCMB28 (D). Human nuclei stain to identify human tumor cells, GPC2 staining for tumor identification and expression, CD3 stain to identify CAR T cells. Scale bar is 2000μm at low power, 250μm at insert. (**D**) Overall survival of mice implanted with thalamic DMG xenograft 7316-3058 treated with six repeated doses of 2 x 10^6^ CAR T cells. Doses indicated by arrows on graph. N= 7 nice per arm. ****, p<0.0001; **, p<0.01; *, p<0.05; ns, not significant.

As both the 7316-4509 and RCMB28 medulloblastoma xenograft models developed clinical signs of xenogeneic GVHD from repeated doses of human T cells prior to tumor end point, we next utilized the clinically aggressive, fast-growing thalamic DMG 7316-3058 xenograft model to evaluate the ability of mRNA GPC2-directed CARs to induce prolonged murine survival. Prior studies treating thalamic DMG xenograft murine models with GD2-directed CAR T cells have resulted in fatal pseudoprogression and hydrocephalus(*35*). However, we hypothesized that serial low doses of GPC2 mRNA CAR T cells might avoid CNS toxicity due to their transient nature, but still provide tumor control. Using lower doses of 2 x 10^6^ GPC2- or CD19-directed CAR T cells also allowed study observations up to 60 days post initial T cell infusion. CAR T cells were delivered intratumorally via catheter for a total of 6 doses, and D3V3 and D3V4 GPC2-directed CAR T cells both significantly prolonged murine survival compared to CD19 CAR T cell controls (median survival of 55 days for D3V3 GPC2 CARs and >60 days for D3V4 GPC2 CARs versus 33 days for CD19 CARs; p<0.05 and p<0.01, respectively) (**Fig 6D**). Again, no toxicity was observed during the prolonged course of this study with these murine GPC2 cross-reacting CAR T cells.

## DISCUSSION

Pediatric brain tumors are the leading cause of death in pediatric oncology (*36*). CAR T cells have transformed the treatment of relapsed and refractory leukemia and lymphoma, with continued pursuit of similar efficacy in solid and brain tumors. Our group recently identified GPC2 as an immunotherapeutic target in neuroblastoma with expression in other pediatric and adult tumors, demonstrating robust and sustained efficacy of a GPC2 ADC utilizing the GPC2 D3 antibody(*25, 26*). We and others have also shown that lentiviral-based GPC2 DNA CARs can lead to significant efficacy in neuroblastoma murine models utilizing alternative GPC2 scFvs(*27, 28, 37*). Here we focus on GPC2 in pediatric brain tumors, both in quantifying GPC2 expression across a wide-range of unique brain tumor histotypes, and on developing mRNA-based GPC2-directed CARs to enable pediatric brain tumor treatment approaches. We describe the creation and manipulation of a CAR construct using the same D3 binding domain utilized in our prior GPC2 ADC work, to optimize performance in CAR T cells.

Investigation of GPC2 expression using RNA sequencing and various proteomic approaches revealed elevated GPC2 expression across diverse groups of pediatric brain tumors. While many of these pediatric brain tumors have embryonal origins similar to neuroblastoma, such as medulloblastoma, ETMR, CNS embryonal tumors NOS, and pineoblastomas, several others do not share this phenotype, including HGG and CPC. H3 G35 mutant tumors showed the highest *GPC2* expression among HGGs, and have been shown to be driven by overexpression of MYCN(*38*). CPC was also recently shown to be driven by activation of MYC paired with loss of TP53(*39*). This increase in MYCN and MYC activity may account for the increased *GPC2* expression in these groups, as we have previously shown MYC family proteins induce higher *GPC2* expression in neuroblastoma(*25*). Across the CBTN cohort, *MYCN* expression was positively correlated with *GPC2* expression, although not in medulloblastoma. Increased *GPC2* expression in medulloblastoma may be driven in part by 7q gain(*25*). *MYC* expression may also play a role, and future work will focus on understanding factors driving *GPC2* expression in all pediatric brain tumors, and identification of other drivers that may be contributing. Our data also show that GPC2 in HGG cell lines seems to coincide with a highly malignant stem-like population of cells and enhanced GPC2 expression could be selected for by using stem cell conditions in cell line development. This is consistent with our recent finding that GPC2 expression is increased in the stem cell compartment of neuroblastomas and small cell lung cancers(*26*), and future work will evaluate the potential oncogenic and stem-like properties of these cells in pediatric brain tumors. Our RNA sequencing data also showed no significant difference in *GPC2* expression across diagnosis-relapse paired samples, suggesting GPC2-directed CARs may be clinically efficacious in both settings.

Using mRNA, we were able to develop multiple CAR T cell constructs from one scFv domain, D3, which was previously used successfully to create an ADC. Our data showed the light-to-heavy configuration of the scFv proved superior, with efficacy *in vivo* using both the 15 amino acid and 5 amino acid linker between the chains. This work was carried out using an mRNA CAR construct for two reasons. First, mRNA is more facile for quick iterative testing of different constructs. Unfortunately, evaluation of optimal configuration for CAR T cells remains empirical, with different targets requiring different adjustments for optimal efficacy. Our mRNA methodology(*14*) allowed for quick testing and prioritization of the D3V3 and D3V4 constructs, and should be utilized for screening of other novel CAR constructs. Second, mRNA CAR T cells may mitigate any potential off-target toxicity when GPC2 CAR T cells are delivered to the brain given their transient CAR expression. In our aggressive thalamic DMG model, we were able to avoid fatal pseudoprogression as has been previously observed in other studies using viral-based CAR T cells targeting thalamic DMG(*35*), by employing serial low doses of mRNA GPC2-directed CAR T cells.

As previously described, mRNA CAR T cells have not been shown to effectively eradicate solid tumors via systemic intravenous injection due to their transient nature(*20*). However, intratumoral or local delivery can produce full tumor regression(*20, 23, 24, 40*), which we have also shown here with our experiments using mRNA GPC2-directed CAR T cells in two medulloblastoma models and one thalamic DMG model. With growing evidence for the effectiveness of locoregional delivery of CAR T cells for brain tumors both preclinically(*32, 41, 42*) and in trials(*9*), the cannula system we developed for repeated CAR T cell delivery serves as an important platform for preclinical CAR T cell testing. The cannulas allow for CAR T cell administration that bypasses the blood brain barrier, similar to intraventricular reservoirs or convection enhanced delivery catheters, with the former currently being used in ongoing clinical trials for CAR T cells in pediatric brain tumors (NCT03500991, NCT03638167, NCT04185038, NCT04196413) (*11*), propelling the translational applicability of this work and providing a clinical platform for future studies in humans.

In conclusion, this work has shown that the immunotherapeutic target GPC2 is highly expressed across a variety of pediatric brain tumors, including malignant embryonal tumors (CNS embryonal tumors NOS, ETMRs, medulloblastomas) and a subset of HGGs and DMGs. Using mRNA we created two efficacious CAR constructs from the D3 binder and show efficacy in several pediatric brain tumor preclinical models, laying the foundation for the translation of GPC2 CARs to pediatric brain tumors.

## MATERIALS AND METHODS

### RNA sequencing of primary tumor samples and patient derived xenografts

RNA sequencing of pediatric brain tumor samples was completed as described within the OpenPBTA(*43*) (available at https://alexslemonade.github.io/OpenPBTA-manuscript/v/0d99c02483e21557e27957386d419b58fe8cb635/ or from the public CAVATICA project: https://cavatica.sbgenomics.com/u/cavatica/openpbta). Briefly, 1,028 paired-end RNA fastq files, comprising 970 rRNA-depleted and 58 poly-A enriched samples, were aligned to the hg38 human genome using GENCODE version 27 as reference annotation with STAR v2.6.1d. Gene level quantification in terms of TPM (transcripts per million) was quantified using RSEM version 1.3.1. The rRNA depleted and poly-A enriched samples were batch corrected using the ComBat function of R package sva.

### Tumor Cell lines

Neuroblastoma cell lines were maintained in culture as previously described(*25, 44*) with RPMI media supplemented with Penicillin/Streptomycin, L-glutamine, and 10% FBS. Brain tumor cell lines were maintained in serum free culture, using DMEM:F12 media supplemented with EGF, FGF and B27. Luciferase was introduced into pediatric brain tumor cell lines using lentiviral plasmids: pLenti CMV Puro LUC (w168-1) was a gift from Eric Campeau & Paul Kaufman(*45*) (Addgene plasmid # 17477; http://n2t.net/addgene:17477; RRID:Addgene_17477) and pLL-EF1a-rFLuc-T2A-GFP-mPGK-Puro (LL410PA-1, System Biosciences). Preparation and transduction was completed as previously described,(*25*) or with pre-made lentivirus expressing firefly luciferase with GFP (PLV-10173-200, Cellomics). Cell lines were tested for mycoplasma and confirmed with STR sequencing.

### T cell expansion and transfection

Human T cells from healthy de-identified donors were purchased from the University of Pennsylvania Immunology Core and stimulated with CD3/CD28 beads (11131D, Gibco) as previously described(*46*). Stimulated and rested T cells were frozen in liquid nitrogen prior to use. After thawing and resting for one day, transfection was completed using Amaxa Nucleofector (Lonza) or ECM 830 (BTX) electroporation systems as previously described(*14*).

### CAR construct cloning

Sequences encoding the D3 scFV were combined with sequences encoding a CD8 hinge and transmembrane domain, followed by 41BB and CD3ζ co-stimulatory domains, to create the CAR constructs. CAR plasmids were generated by GenScript, and then cloned into a pTEV vector backbone using Geneart Seamless cloning (A14603, Invitrogen).

### In vitro mRNA synthesis

mRNA encoding CAR constructs were synthesized from the pTEV vectors using Megascript T7 transciption kit (AM1334, Invitrogen) and replacing UTP with 1-methylpseudouridine triphosphate (N-1081, TriLink Biotechnologies) as previously described(*14*). After synthesis, all mRNA was purified with RNase III (RN02950, Epicenter), followed by capping and tailing as previously described(*14*).

### Western blot validation of protein expression

Tumor cell lines were lysed and western blotting completed as previously described(*25*), using anti-Glypican-2 (1:500; F-5, sc-393824, Santa Cruz Biotechnology) for protein detection.

### Flow cytometry

Flow cytometry was performed on tumor cell lines to confirm GPC2 expression on the cell surface. Tumor cell lines were washed in FACS buffer (500 mL PBS, 10 mL FBS, 2 mL 0.5 M EDTA), then incubated in D3-GPC2-IgG1 conjugated to phycoerythrin (PE) at 1:800 for 20 minutes in the dark at 4° C. Conjugation with PE was completed using conjugation kit (ab102918, Abcam). For quantification of surface GPC2, BD Quantibrite beads (340495, BD Biosciences) were used per manufacturer’s instruction.

CAR T cell constructs were evaluated for GPC2 specific binding by staining with GPC2 protein (R&D) conjugated to PE. CAR T cells were washed in FACS buffer and incubated in GPC2-PE at 1:100 for 20 minutes in the dark at 4° C, and then washed again in FACS buffer before analysis. CAR T cells were evaluated for CAR expression on the surface by flow cytometry with protein L (1:100; M00097, Genscript) as previously described(*47*). Flow cytometry for negative checkpoint regulator was performed as previously described(*14*) using antibodies: BV510 mouse anti-human CD279 (PD1) (1:100; 563076, BD Horizon), PerCP-eFluor 710 mouse anti-human CD223 (LAG-3) (1:100; 46-2239-41, Invitrogen), and BV421 mouse anti-human CD366 (TIM-3) (1:100; 565562, BD Biosciences).

Flow cytometry data was acquired on BD Accuri C6 (BD Biosciences) or FACS Verse (BD Biosciences) flow cytometers. Analysis was completed on FlowJo 10.2 (Treestar).

### Cytotoxicity assays

Measurement of CAR T cell dependent tumor cell killing was measured by several methods. For adherent tumor cell lines, cell proliferation was measured using Real-time Excelligence system (RT-CES, F Hoffman La-Roche) measured every hour. For suspension cell lines, luciferase containing tumor cells were plated in 96 well plate, CAR T cells plated after 24 hours, and Bright Glo luciferase assay (E2620, Promega) was run after 48 hours of co-incubation. CAR T cells were plated in at least triplicate on tumor cells at E:T ratios of 10:1, 5:1 and 1:1. Each experiment was repeated at least two times.

### CAR T Cell cytokine release

Tumor specific CAR T cell degranulation was determined using ELISA to detect interferon-gamma and IL-2. Tumor cells and CAR T cells were plated as described above and allowed to co-incubate for 24-48 hours. Plates were then spun and supernatant collected for evaluation using human IL-2 duoset ELISA (DY202, R&D) and human IFN-gamma duoset ELISA (D285B, R&D) per manufacturer’s instructions.

### Mouse studies

*In vivo* studies were completed in mice under an approved protocol by the Institutional Animal Care and Use Committee of the Children’s Hospital of Philadelphia (IAC 19-000907). Studies used 6- to 10-week old NOD-SCID-γc–/– (NSG) mice obtained from Jackson Laboratories or bred in-house under pathogen free conditions.

For medulloblastoma studies, RCMB28 xenograft tumors were provided by Dr. Robert Wechsler-Reya, and 7316-4509 provided by the CBTN. Medulloblastoma xenografts were implanted stereotactically into the cerebellum of mice using the coordinates from lambda of: A/P -2.00mm, D/V 2.00mm, M/L -2.00mm. For thalamic DMG model, 7316-3058 was provided by the CBTN. This model was injected into the thalamus using the coordinates from bregma: A/P - 1.00mm, D/V 3.50mm, M/L -0.80mm. Following engraftment, indwelling catheters to deliver intratumoral CAR T cells in the CNS were introduced using 26 gauge guide cannulas (C315GS-5/SPC, P1 Technologies), stereotactically placed and secured to the skull using screws (C212SG, P1 Technologies) and acrylic resin (Ortho-Jet, Lang). Catheters were placed at the same coordinates as the tumor cell placement. T cells were infused using 33 gauge internal cannulas (C315IS-5/SPC, P1 Technologies) placed into the guide cannula and delivered over 1 minute in a Hamilton syringe. All studies included 6-10 animals per arm.

### Immunohistochemical analysis

Primary tumor samples, tissue microarrays and xenograft tumors from sacrificed animals were stained for GPC2 using immunohistochemistry as previously described,(*25*) by the Children’s Hospital of Philadelphia Pathology Core laboratory. Staining for GPC2 was graded in a blinded fashion by a neuropathologist. Animals treated with CAR T cells had tumors excised and were also stained with H&E and human CD3 (A0452, Dako), in addition to GPC2. Photographs were taken using a Leica DM4000B.

### Statistical analysis

All data generated was evaluated using Prism 8 (Graphpad). Comparison of groups for *GPC2* expression was completed using two-sided Mann-Whitney test or Kruskal-Wallis test. Correlation was calculated using two-tailed Pearson. Comparison of means for *in vitro* and *in vivo* CAR T cell data was completed using two-sided student t-test and 2-way ANOVA. P-values <0.05 were considered statistically significant. Data is generally presented as the mean ± SEM or as noted in the figure and figure legend. Number of samples or replicates is indicated in the results, figures and figure legends.

## Acknowledgments

This research was conducted using data and samples made available by the Children’s Brain Tumor Network. We thank the patients and families who generously donated their tumor specimens.

## Funding

Alex’s Lemonade Stand Foundation (K.R.B.)

Children’s Hospital of Philadelphia, Giulio D’Angio Endowed Chair (J.M.M.)

Damon Runyon Cancer Research Foundation PST-07-16 (K.R.B.)

EVAN Foundation (K.R.B.) Grayson Saves Foundation (J.B.F.)

Hyundai Hope on Wheels Young Investigator Award (J.B.F)

Kortney Rose Foundation (J.B.F.)

National Institutes of Health NCI K12 CA076931-19 (J.B.F.)

National Institutes of Health NCI K08 CA230223 (K.R.B.)

National Institutes of Health NCI U54 CA232568 (J.M.M.)

National Institutes of Health NCI R35 CA220500 (J.M.M.)

St. Baldrick’s-Stand Up to Cancer Dream Team Translational Research Grant SU2C-AACR-DT-27-17 (J.M.M.). The St. Baldrick’s Foundation collaborates with Stand Up To Cancer. Research Grants are administered by the American Association for Cancer Research, the Scientific Partner of SU2C.

## Author contributions

Conceptualization: J.B.F., D.M.B., A.C.R., J.M.M., and K.R.B.

Methodology: J.B.F., T.S., D.M.B., and K.R.B.

Software, J.L.R. and K.R.

Data Curation: J.L.R. and K.R.

Formal analysis: J.B.F.

Investigation: J.B.F., C.G., A.S., C.B., M.L, S.B., P.M., and D.M.

Resources: R.W.R., K.K., A.C.R., J.M.M., and K.R.B.

Supervision: D.M.B., A.C.R., J.M.M., and K.R.B.

Funding Acquisition: J.B.F., P.B.S., A.C.R., J.M.M., and K.R.B.

Writing – Original Draft: J.B.F.

Writing – Review & Editing: J.B.F., C.G., J.L.R., A.S., C.B., K.R., M.V.L., S.N.B., T. S., P.J.M., D.M., R.J.W., K.K., P.B.S., D.M.B, A.C.R., J.M.M., and K.R.B.

## Data and materials availability

All data are available in the main text.

## References and Notes

1. J. Couzin-Frankel, Cancer Immunotherapy. Science 342, 1432–1433 (2013).

2. H. Almasbak, T. Aarvak, M. C. Vemuri, CAR T Cell Therapy: A Game Changer in Cancer Treatment. J Immunol Res 2016, 5474602 (2016).

3. S. L. Maude et al., Chimeric antigen receptor T cells for sustained remissions in leukemia. N Engl J Med 371, 1507–1517 (2014).

4. T. J. Fry et al., CD22-targeted CAR T cells induce remission in B-ALL that is naive or resistant to CD19-targeted CAR immunotherapy. Nature Medicine 24, 20 (2017).

5. R. Gardner et al., CD19CAR T Cell Products of Defined CD4:CD8 Composition and Transgene Expression Show Prolonged Persistence and Durable MRD-Negative Remission in Pediatric and Young Adult B-Cell ALL. Blood 128, 219–219 (2016).

6. First-Ever CAR T-cell Therapy Approved in U.S. Cancer Discovery 7, OF1–OF1 (2017).

7. J. Wagner, E. Wickman, C. DeRenzo, S. Gottschalk, CAR T Cell Therapy for Solid Tumors: Bright Future or Dark Reality? Mol Ther 28, 2320–2339 (2020).

8. D. M. O’Rourke et al., A single dose of peripherally infused EGFRvIII-directed CAR T cells mediates antigen loss and induces adaptive resistance in patients with recurrent glioblastoma. Science Translational Medicine 9, eaaa0984 (2017).

9. C. E. Brown et al., Regression of Glioblastoma after Chimeric Antigen Receptor T-Cell Therapy. N Engl J Med 375, 2561–2569 (2016).

10. C. E. Brown et al., Bioactivity and Safety of IL13Rα Receptor CD8^+^ T Cells in Patients with Recurrent Glioblastoma. Clinical Cancer Research 21, 4062–4072 (2015).

11. N. A. Vitanza et al., Locoregional infusion of HER2-specific CAR T cells in children and young adults with recurrent or refractory CNS tumors: an interim analysis. Nat Med, (2021).

12. S. Feins, W. Kong, E. F. Williams, M. C. Milone, J. A. Fraietta, An introduction to chimeric antigen receptor (CAR) T-cell immunotherapy for human cancer. American Journal of Hematology 94, S3–S9 (2019).

13. J. B. Foster, D. M. Barrett, K. Karikó, The Emerging Role of In Vitro-Transcribed mRNA in Adoptive T Cell Immunotherapy. Molecular Therapy 27, 747–756 (2019).

14. J. B. Foster et al., Purification of mRNA Encoding Chimeric Antigen Receptor Is Critical for Generation of a Robust T-Cell Response. Hum Gene Ther 30, 168–178 (2019).

15. N. Pardi, M. J. Hogan, D. Weissman, Recent advances in mRNA vaccine technology. Current Opinion in Immunology 65, 14–20 (2020).

16. D. M. Barrett et al., Treatment of advanced leukemia in mice with mRNA engineered T cells. Hum Gene Ther 22, 1575–1586 (2011).

17. N. Singh, D. M. Barrett, S. A. Grupp, Roadblocks to success for RNA CARs in solid tumors. Oncoimmunology 3, e962974 (2014).

18. R. A. Morgan et al., Cancer regression and neurological toxicity following anti-MAGE-A3 TCR gene therapy. J Immunother 36, 133–151 (2013).

19. J. Carroll, Exclusive: Carl June’s Tmunity encounters a lethal roadblock as 2 patient deaths derail lead trial, raise red flag forcing rethink of CAR-T for solid tumors. Endpoints News. 2021.

20. N. Singh et al., Nature of tumor control by permanently and transiently modified GD2 chimeric antigen receptor T cells in xenograft models of neuroblastoma. Cancer Immunol Res 2, 1059–1070 (2014).

21. X. Liu et al., Novel T cells with improved in vivo anti-tumor activity generated by RNA electroporation. Protein & Cell 8, 514–526 (2017).

22. G. L. Beatty et al., Activity of Mesothelin-Specific Chimeric Antigen Receptor T Cells Against Pancreatic Carcinoma Metastases in a Phase 1 Trial. Gastroenterology 155, 29–32 (2018).

23. W. X. Ang et al., Intraperitoneal immunotherapy with T cells stably and transiently expressing anti-EpCAM CAR in xenograft models of peritoneal carcinomatosis. Oncotarget 8, 13545–13559 (2017).

24. G. L. Beatty et al., Mesothelin-specific chimeric antigen receptor mRNA-engineered T cells induce anti-tumor activity in solid malignancies. Cancer Immunol Res 2, 112–120 (2014).

25. K. R. Bosse et al., Identification of GPC2 as an Oncoprotein and Candidate Immunotherapeutic Target in High-Risk Neuroblastoma. Cancer Cell 32, 295–309 e212 (2017).

26. S. Raman et al., A GPC2 antibody-drug conjugate is efficacious against neuroblastoma and small-cell lung cancer via binding a conformational epitope. Cell Reports Medicine 2, 100344 (2021).

27. N. Li, H. Fu, S. M. Hewitt, D. S. Dimitrov, M. Ho, Therapeutically targeting glypican-2 via single-domain antibody-based chimeric antigen receptors and immunotoxins in neuroblastoma. Proc Natl Acad Sci U S A 114, E6623–E6631 (2017).

28. N. Li et al., CAR T cells targeting tumor-associated exons of glypican 2 regress neuroblastoma in mice. Cell Reports Medicine 2, 100297 (2021).

29. H. Ijaz et al., Pediatric high-grade glioma resources from the Children’s Brain Tumor Tissue Consortium. Neuro Oncol 22, 163–165 (2020).

30. A. Wenger et al., Stem cell cultures derived from pediatric brain tumors accurately model the originating tumors. Oncotarget 8, (2017).

31. E. O. Vik-Mo et al., Brain tumor stem cells maintain overall phenotype and tumorigenicity after in vitro culturing in serum-free conditions. Neuro-Oncology 12, 1220–1230 (2010).

32. J. Theruvath et al., Locoregionally administered B7-H3-targeted CAR T cells for treatment of atypical teratoid/rhabdoid tumors. Nature Medicine, (2020).

33. J. M. Rusert et al., Functional Precision Medicine Identifies New Therapeutic Candidates for Medulloblastoma. Cancer Res 80, 5393–5407 (2020).

34. D. M. Barrett et al., Regimen-specific effects of RNA-modified chimeric antigen receptor T cells in mice with advanced leukemia. Hum Gene Ther 24, 717–727 (2013).

35. C. W. Mount et al., Potent antitumor efficacy of anti-GD2 CAR T cells in H3-K27M(+) diffuse midline gliomas. Nat Med 24, 572–579 (2018).

36. S. C. Curtin, A. M. Minino, R. N. Anderson, Declines in Cancer Death Rates Among Children and Adolescents in the United States, 1999-2014. NCHS Data Brief, 1–8 (2016).

37. S. Heitzeneder et al., Abstract A09: Glypican-2 targeted CAR T cells designed to effectively eradicate endogenous site density solid tumors in the absence of toxicity. Cancer Research 80, A09–A09 (2020).

38. L. Bjerke et al., Histone H3.3 Mutations Drive Pediatric Glioblastoma through Upregulation of MYCN. Cancer Discovery 3, 512–519 (2013).

39. J. Wang et al., Myc and Loss of p53 Cooperate to Drive Formation of Choroid Plexus Carcinoma. Cancer Research 79, 2208–2219 (2019).

40. K. Schutsky et al., Rigorous optimization and validation of potent RNA CAR T cell therapy for the treatment of common epithelial cancers expressing folate receptor. Oncotarget 6, 28911–28928 (2015).

41. L. K. Donovan et al., Locoregional delivery of CAR T cells to the cerebrospinal fluid for treatment of metastatic medulloblastoma and ependymoma. Nature Medicine, (2020).

42. C. E. Brown et al., Optimization of IL13Rα Cells for Improved Anti-tumor Efficacy against Glioblastoma. Molecular Therapy 26, 31–44 (2018).

43. S. C. Shapiro JA, Bethell CJ, Gaonkar KS, Zhu Y, Brown MA, et al., An open pediatric brain tumor atlas. Manubot [Internet], https://alexslemonade.github.io/OpenPBTA-manuscript/v/7207b5942e7207c7205ee7208a7363f7202cc7254c7204a7278ec7206f7810e/ (2020).

44. J. L. Harenza et al., Transcriptomic profiling of 39 commonly-used neuroblastoma cell lines. Scientific Data 4, 170033 (2017).

45. E. Campeau et al., A versatile viral system for expression and depletion of proteins in mammalian cells. PLoS One 4, e6529 (2009).

46. C. Carpenito et al., Control of large, established tumor xenografts with genetically retargeted human T cells containing CD28 and CD137 domains. Proc Natl Acad Sci U S A 106, 3360–3365 (2009).

47. Z. Zheng, N. Chinnasamy, R. A. Morgan, Protein L: a novel reagent for the detection of Chimeric Antigen Receptor (CAR) expression by flow cytometry. Journal of Translational Medicine 10, 29 (2012).

